# Elevated L1 expression in ataxia telangiectasia likely explained by an RNA-seq batch effect

**DOI:** 10.1101/2022.10.06.511227

**Authors:** Geoffrey J. Faulkner

**Affiliations:** Mater Research Institute - University of Queensland, Woolloongabba, QLD, 4102, Australia; Queensland Brain Institute, University of Queensland, St. Lucia, QLD, 4067, Australia

## Abstract

A recent study (Takahashi et al., *Neuron*, 2022) concluded LINE-1 (L1) retrotransposon activation drives cerebellar ataxia and neurodegeneration. This position was based on L1 upregulation in ataxia telangiectasia (AT) patient cerebellum samples, as measured by RNA-seq, and observation of ataxia and neurodegeneration in mice where cerebellar L1 expression was induced via dCas9-CRISPR. Here, a re-analysis of the RNA-seq data, which were obtained by rRNA depletion rather than polyA+ selection, revealed a high fraction (38.4%) of intronic reads. Significantly (*p*=0.034) more intronic reads were present in the AT data than the matched controls. This finding provides an alternative and robust explanation for a key result reported by Takahashi et al.: intronic L1 sequences are abundant in pre-mRNAs, and more pre-mRNAs were retained in the AT libraries. This apparent batch effect deserves further examination, as claims of L1-mediated pathogenesis could shape future efforts to treat AT by trying to attenuate L1 activity.

## Main

The retrotransposon LINE-1 (L1) is an autonomous mobile genetic element whose ~500,000 copies compose more than 17% of the human genome. A full-length, retrotransposition-competent L1 is 6kb long and incorporates a 5′ internal sense promoter that drives transcription of a bicistronic mRNA, which in turn encodes two proteins (ORF1p and ORF2p). These L1 proteins strongly prefer to mobilize their encoding mRNA, and there are fewer than 100 retrotransposition-competent human-specific L1 (L1HS) copies per individual. The vast majority of L1s scattered throughout the genome thus do not have intact ORFs and are immobile (Kazazian and Moran, 2017), even if they reside in the introns of protein-coding genes.

Ataxia telangiectasia (AT) is a severe neurodegenerative disorder caused by defects in the AT mutated (ATM) DNA damage repair gene. Prior works have found ATM either facilitates L1 mobilization (Gasior et al., 2006; Wallace et al., 2013) or, if mutated, changes the character and, moderately, increases the frequency of L1 insertions found in neuronal cells (Coufal et al., 2011). ATM is not however thought to repress L1 promoter activity or limit L1 protein expression (Coufal et al., 2011).

In recent work, Takahashi et al. concluded that L1 can drive neurodegeneration in AT, based on increased cerebellar L1 RNA abundance in AT patients, as well as ataxia and neurodegeneration in mice where L1 transcription was activated in the cerebellum via a dCas9-CRISPR approach (Takahashi et al., 2022). This intriguing study could shape strategies to treat or prevent AT in children and, for this reason, its findings warrant careful consideration.

Takahashi et al. used bulk RNA-seq, where libraries were prepared via rRNA depletion, to compare the cerebellar transcriptomes of AT patients (*n*=6) and control individuals (*n*=6). They reported significant upregulation of the L1HS subfamily in AT. Accurate quantification of L1HS transcription, and that of other recently emerged, or young, L1 subfamilies with RNA-seq presents many challenges, principally due to their copy number and limited sequence divergence (Faulkner et al., 2009; Iñiguez et al., 2019; Jin et al., 2015; Lanciano and Cristofari, 2020). When evaluating differential expression amongst numerous L1 subfamilies, Takahashi et al. did not incorporate a multiple testing correction despite performing t tests on several retrotransposon families simultaneously, which would have rendered the noted L1HS upregulation in AT patients (*p*=0.042, two-sided t test) no longer significant. Multiple testing correction and the incorporation of reads that align to multiple genomic locations, or “multi-mapping” reads are essential to the robust and reproducible RNA-seq quantification of young retrotransposon subfamilies, such as L1HS (Jin et al., 2015; Lanciano and Cristofari, 2020).

Re-analyzing the RNA-seq data with TEtranscripts (Jin et al., 2015) and the same STAR (Dobin et al., 2013) alignment parameters used by Takahashi et al., a statistically significant difference was not apparent for L1HS in AT patients compared to controls, and incorporating multi-mapping reads did not change this outcome (**Figure S1A**). Older L1 subfamilies such as L1ME, which are immobile and tend to be 5′ truncated and lack a canonical promoter and intact ORFs, accounted for far more RNA-seq reads than L1HS (**Figure S1A**). These older L1s were also typically more highly expressed in the AT samples than in controls, as were young PolIII-transcribed *Alu* retrotransposon families, such as AluYa5 (**Figure S1A**). Visual inspection of a selection of the most highly expressed L1 loci, such as an L1HS intronic to the *TSPAN5* gene (**Figure S1C**) and an L1ME intronic to the *DPP6* gene (**Figure S1C**), indicated they did not possess intact ORFs and were typically flanked by pronounced and pervasive intronic RNA-seq signals (**Figure S1C**).

RNA-seq libraries prepared via rRNA depletion, as opposed to polyA+ selection, can contain considerably more nascent, unspliced pre-mRNAs (Zhao et al., 2018). While the average percentage of intragenic reads did not appreciably differ in control (87.6%) and AT samples (88.3%), the average percentage of intronic reads was both unusually high overall (38.4%) and significantly (*p*=0.034, two-tailed t test with Bonferroni correction) higher in AT samples (45.2%) than in control samples (31.6%) (**Figure S1B**). Normalized L1HS, L1ME and AluYa5 read counts were each strongly correlated (Pearson *r* >0.92, two-tailed *p*<0.01) with intronic read fraction in both AT and control samples (**Figure S1D**). Further normalization by dividing the L1HS read count by the fraction of intronic reads detected in each sample brought L1HS expression to virtual parity in AT and control samples (**Figure S1E**). The expression of L1HS repressors downregulated in AT, as noted by Takahashi et al., was also strongly anticorrelated with intronic read fraction, such as for *TRIM28* (Pearson *r*=-0.90, two-tailed *p*<0.0001), which has a relatively high (1:1) exon:intron sequence ratio and, notably, regulates older L1 subfamilies and not L1HS transcription (Castro-Diaz et al., 2014). Despite the availability of robust human L1 ORF1p antibodies, an immunoblot was not shown to support differential L1 protein expression in AT patient samples.

To corroborate the RNA-seq results, qPCR was performed. However, this approach depends on high quality input RNA and, as the target L1HS RNA is not spliced, is prone to gDNA contamination. These factors, and off-target effects, can influence L1 qPCR assays, particularly when primers are designed against the L1HS ORFs, as done here, and not the 5′UTR, which is retained only by full-length L1s. Finally, to demonstrate the robustness of the present claims, normalized L1HS read counts were obtained by aligning the RNA-seq data with bowtie2 (Langmead and Salzberg, 2012) and then analyzed with RepEnrich2 (Criscione et al., 2014), as per the approach of Takahashi et al., confirming a strong correlation (Pearson *r*=0.97, two-tailed *p*<0.0001) with intronic read fraction across AT and control samples (**Figure S1F**). This analysis confirmed the strong correlation of L1HS and intronic read count remained, regardless of the alignment algorithm, TE transcript analysis software, or how exonic/intronic regions were defined (see **Methods**).

**Figure S1:**
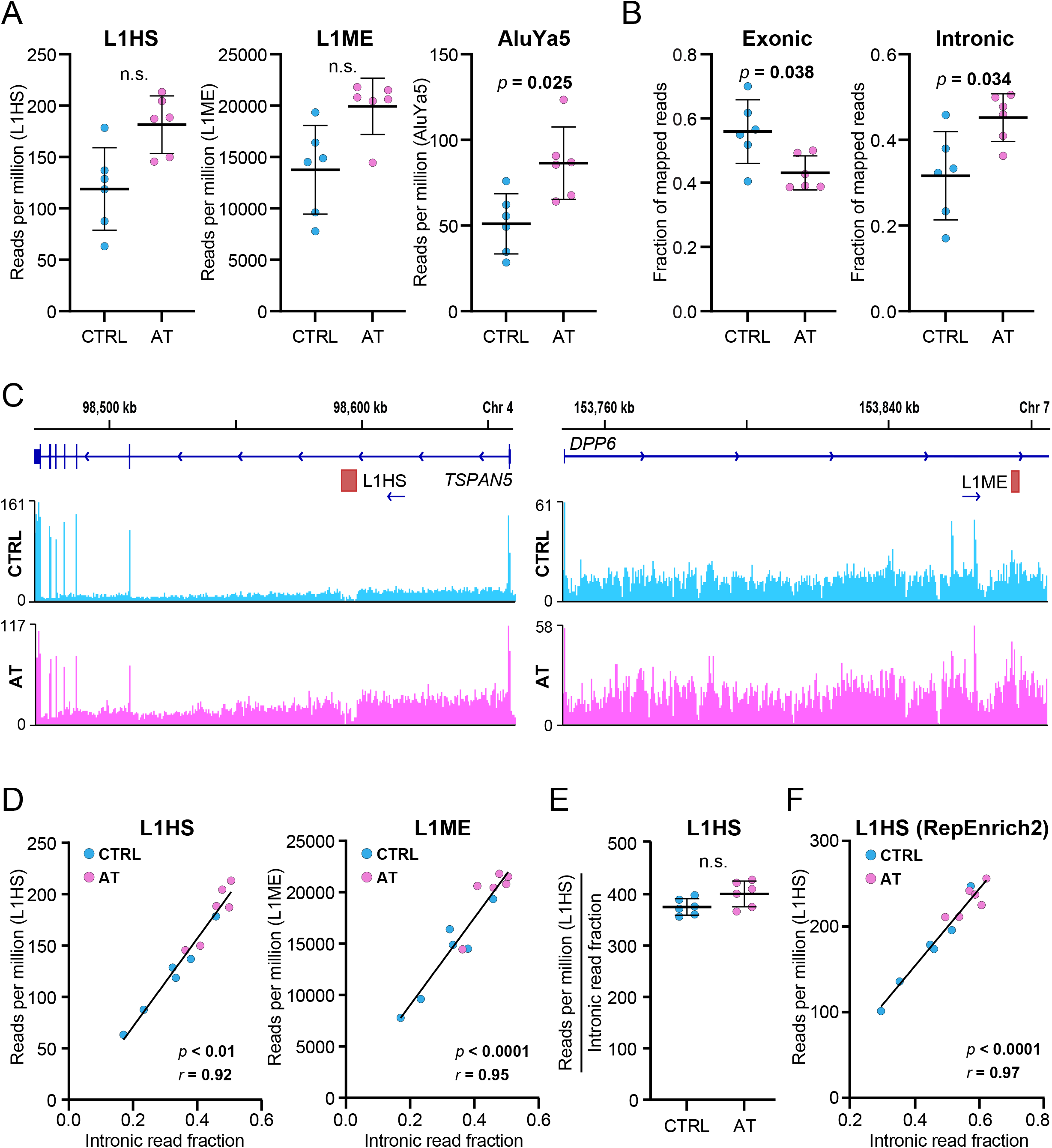
Intronic enrichment from retained pre-mRNAs in AT patient RNA-seq data explains apparent retrotransposon upregulation. **(A)** Normalized RNA-seq read counts in retrotransposon subfamilies obtained from TEtranscripts and allowing multi-mapping reads. Statistical significance was calculated using DESeq2 and BH multiple testing correction. **(B)** Fractions of exonic and intronic reads per sample, using only uniquely mapping reads, determined with featurecounts. Significance testing was via two-tailed t test with Bonferroni correction. **(C)** Integrative Genomics Viewer (IGV) view of aggregate RNA-seq coverage plots at the *TSPAN5* locus (left) and *DPP6* locus (right). Each L1 has broken ORFs in the annotated reference genome sequence and is oriented on the same strand as the host gene. **(D)** Correlation of L1 subfamily normalized read count, obtained with TEtranscripts. and intronic read fraction per sample, with a linear regression line and Pearson correlation (*r* and two-tailed *p*) shown. **(E)** Normalized L1HS read count divided by intronic read fraction per sample. **(F)** As per (D), except showing normalized L1HS read counts obtained using RepEnrich2 and intronic read fractions determined using the bowtie2 alignment input files for RepEnrich2 via rnaseqc. Note: in (A), (B) and (E), individual data points, each representing a sample, are marked, and also represented as mean ± S.D.

Taken together, these results point to a technical explanation (Zhao et al., 2018) for increased L1 transcript abundance in AT patients: pre-mRNAs, whose introns are substantially composed of L1 and other retrotransposons, were more abundant in the AT sample RNA-seq libraries than in the matched control samples. The most likely explanation for this difference is a batch effect where the RNA quality was lower for the AT samples than for the control samples, or perhaps an effect arising during the rRNA depletion-based library processing. Ideally, the authors could provide further information distinguishing these possibilities. A much less plausible explanation is that pre-mRNAs are for unknown reasons more abundant in AT cerebellum than in controls, and even this scenario does not yield support for elevated protein-coding L1 transcription in AT.

A finding of L1 not being upregulated in AT would be concordant with earlier data suggesting the L1HS promoter is not more active, and L1 proteins not more abundant, in ATM-deficient human neuronal precursor cells (Coufal et al., 2011). It would subtract the human disease rationale for the remainder of the Takahashi et al. study, contradict the model of AT pathology being potentially driven by elevated and ectopic L1-mediated reverse transcription, and, importantly, argue against the therapeutic potential of L1 reverse transcriptase inhibitors in AT.

Even if the available evidence of L1 dysregulation in human AT is discounted, the animal experiments conducted by Takahashi et al. provide a remarkable proof-of-principle of L1 activation *in vivo* via the dCas9-CRISPR system. Previously, AT mouse models have not recapitulated the neurodegeneration seen in AT patients (Lavin, 2013) or the L1 activator dCas9-CRISPR mice generated by Takahashi et al. As a caveat they note, these *in vivo* data were generated using one L1 sgRNA matching thousands of genomic loci, elevating the probability of an off-target effect. This issue is potentially more acute given inadvertent basal dCas9 expression in the L1 activation system, and compounded by the lack of cerebellar RNA-seq data from the dCas9-CRISPR mice, as L1 sequences can act as promoters for genes expressed during development and in neurons (Gerdes et al., 2022; Jönsson et al., 2019). The loci containing the integrated L1 targeting and scrambled sgRNA transgenes were not identified, and hence the effects of their integration, and their genomic context, in combination with the genetic background of the dCas9-CRISPR line, on phenotype was not indicated. While treatment with the L1 reverse transcriptase inhibitor lamivudine appeared to attenuate disease progression in dCas9-CRISPR animals, a relatively low concentration of lamivudine, which crosses the blood-brain barrier very poorly (Osborne et al., 2020), was used. The cerebellar concentration of lamivudine, and whether it was sufficient to alter L1 reverse transcriptase activity, was therefore not determined.

In light of these open questions, more data is needed to ascertain whether L1 expression is dysregulated in AT. Independent replication experiments using RNA-seq protocols that avoid pre-mRNAs, which neither rRNA depletion-based or single-nucleus RNA-seq (snRNA-seq) approaches easily achieve, could be useful in this regard. These data would help clarify the causal role, if any, played by L1 in human neurodegeneration, and indicate its value as a therapeutic target.

## Methods

Human AT patient RNA-seq fastq files (accession GSE175776) were downloaded from the Gene Expression Omnibus and aligned to the hg38 reference genome using STAR (Dobin et al., 2013). Reads were aligned in two ways. Firstly, to retain only uniquely aligned reads, the same STAR alignment parameters as Takahashi et al. were used. Secondly, to incorporate multi-mapping reads, the STAR alignment parameter outFilterMultimapNmax was changed from 1 to 10000. Each set of alignments were then processed with TEtranscripts (Jin et al., 2015) with default parameters, using “uniq” for the first set of alignments and “multi” for the second set of alignments. RefSeq gene and retrotransposon annotation files were obtained from the UCSC Table Browser and the TEtranscripts GitHub, respectively. Differential gene and retrotransposon expression analyses were performed with DESeq2 (Love et al., 2014) and default (BH) multiple testing correction, as implemented as part of TEtranscripts. Exonic and intronic read fractions were obtained using the RefSeq gene annotation file and featureCounts (Liao et al., 2014) with parameters -p -T 8 -B -O. TEtranscript quantification of retrotransposon families included multi-mapping reads; all other analyses used uniquely mapping reads only. For the analysis shown in Figure S1F, normalized L1HS read counts were obtained using RepEnrich2 (Criscione et al., 2014) and bowtie2 (Langmead and Salzberg, 2012), whilst intronic read fractions were determined using the bowtie2 alignment output files and rnaseqc (DeLuca et al., 2012) and Gencode transcript annotations.

## Acknowledgements

The author thanks Dr Sandra Richardson, Dr Adam Ewing, and members of the Faulkner laboratory for helpful discussions. The author receives funding from the Australian NHMRC (GNT1173711) and ARC (DP200102919), Cancer Australia (2003170), and the Mater Foundation.

